# Inferring human population size and separation history from multiple genome sequences

**DOI:** 10.1101/005348

**Authors:** Stephan Schiffels, Richard Durbin

**Author notes:** Correspondence should be addressed to: S.S. or R.D.

## Abstract

The availability of complete human genome sequences from populations across the world has given rise to new population genetic inference methods that explicitly model their ancestral relationship under recombination and mutation. So far, application of these methods to evolutionary history more recent than 20-30 thousand years ago and to population separations has been limited. Here we present a new method that overcomes these shortcomings. The Multiple Sequentially Markovian Coalescent (MSMC) analyses the observed pattern of mutations in multiple individuals, focusing on the first coalescence between any two individuals. Results from applying MSMC to genome sequences from nine populations across the world suggest that the genetic separation of non-African ancestors from African Yoruban ancestors started long before 50,000 years ago, and give information about human population history as recently as 2,000 years ago, including the bottleneck in the peopling of the Americas, and separations within Africa, East Asia and Europe.

---

Human genome sequences are related to each other through common ancestors. Estimates of when these ancestors lived provide insight into ancestral population sizes and ancestral genetic separations as a function of time. In the case of non-recombining loci such as the maternally inherited mtDNA or the paternally inherited Y chromosome, the age of common ancestors can be estimated from the total number of differences between sequences [1–4]. For autosomal sequences, which account for the vast majority of heritable sequence, the reconstruction of genealogical relationships is more complicated due to ancestral recombinations which separate many different genealogical trees in different locations of the genome. While inferring this pattern is more challenging, it provides in principle much more information about our past than non-recombining loci, since only a few samples provide many effectively independent genealogies, allowing inference of an age distribution of common ancestors with high resolution.

Reconstruction of the full underlying ancestral recombination graph is challenging, because the space of possible graphs is extremely large. A substantial simplification was proposed by McVean and Cardin who model the generating process of the genealogical trees as Markovian along the sequence [5, 6]. An application of this idea was presented by Li and Durbin in the Pairwise Sequential Markovian Coalescent (PSMC) model [7], a method which focuses on modeling the two genome sequences in one diploid individual. Because only two sequences are modeled, the coalescent event joining the sequences at the most recent common ancestor is almost always older than 20 thousand years ago (kya), so PSMC can only infer population size estimates beyond 20 kya. Also, with only two sequences there was only limited scope for the analysis of population separations.

For more than two sequences, extending PSMC in the natural way by enumerating all possible trees with their branch lengths along the sequences would be very computationally costly, even under the Markovian model. A recent simplification was suggested by Song and coworkers [8–10], which is based on approximating the conditional sampling process for adding an (*n*+1)th sequence to the distribution of genealogies connecting *n* sequences. In this paper we propose an alternative approach that we call Multiple Sequential Markovian Coalescent (MSMC), which overcomes the increase in complexity by introducing a different simplification. We characterize the relationship at a given location between multiple samples through a much reduced set of variables: i) the time to the most recent common ancestor of *any* two sequences, i.e. the first coalescence, along with the identities of the two sequences participating in the first coalescence (see Figure 1a), and ii) the total length of all singleton branches in the tree, i.e. branches which give rise to variants of minor allele count 1 if hit by a mutation. Given a demographic model we can keep track of the likelihood distribution for these variables based on the observed mutation data as we move along the sequences. As detailed in *Online Methods,* we derive approximate transition and emission rates using the Sequentially Markovian Coalescent (SMC’) [5, 6] framework. This approach allows efficient maximum likelihood estimation of the free parameters, which include the piecewise constant population size as a function of time. If sequences are sampled from different subpopulations, we use additional free parameters for coalescence rates within and across population boundaries, which allow us to infer how subpopulations separated over time. We compare further the conditional sampling approach and our first coalescent approach in the Discussion.

**Figure 1:**
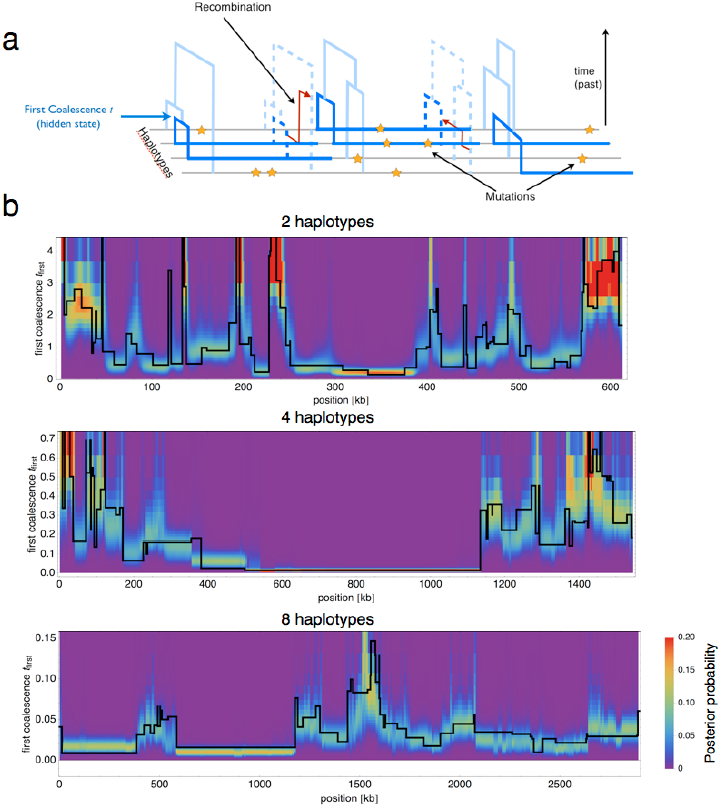
MSMC locally infers branch lengths and coalescence times from observed mutations. (**a**) A schematic representation of the model. Local genealogies change along the sequences by recombination events that rejoin branches of the tree, according to the SMC’ model [5, 6]. The pattern of mutations depends on the genealogy, with few mutations on branches with recent coalescences and more mutations in deeper branches. The hidden states of the model are the time to the first coalescence, along with the identity of the two sequences participating in the first coalescence (dark blue), (**c**) MSMC can locally infer its hidden states, shown by the posterior probability in color shading. In black, we plot the first coalescence time as generated by the simulation. This local inference works well for two, four and eight haplotypes. The more haplotypes, the more recent is the typical time to the first coalescence event, while the typical segment length increases.

Because MSMC focuses on the first coalescence event between any pair of haplotypes, the inference limits are set by the distribution of first coalescence times, the mean of which scales inversely with the square of the sample size, 〈*t*) = 2/(*M*(*M* — 1)) in units of 2*N_0_* generations (see *Online Methods* and Figure 1b). Here we demonstrate application of our model on up to eight haplotypes, which allows us to study population size changes as recent as 70 generations ago. As a special case of MSMC for two haplotypes, we provide a new implementation of PSMC that we call PSMC’ because it uses the SMC’ model, which accounts for recombinations between segments with the same coalescent time [6]. PSMC’ accurately estimates the recombination rate (see *Supplementary* Figure 1), which is not the case for PSMC[7].

We apply our method to 34 individuals from 9 populations of European, Asian, African and Native American ancestry, sequenced using Complete Genomics technology [11]. Our results give detailed estimates of the population sizes and population separations over time between about 2kya and 200kya, including the out-of-Africa dispersal of modern humans, the split between Asian and European populations, and the migration into America.

Like other inference methods based on coalescent theory, MSMC can only infer scaled times and population sizes. To convert them to real time and size, scaled results need to be divided by the mutation rate and scaled times in addition multiplied by the generation time. Here we will use a generation time of 30 years and a rate of 1.25×10^−8^ mutations per nucleotide per generation, as supported by recent publications [12–16]. Since there is current debate about these values [17–19] we consider alternate scalings in the Discussion.

## Results

### Testing the performance of MSMC with simulated data

We implemented the MSMC model as described in *Online Methods,* with mathematical derivations given in *Supplementary Note.* To test the model, we used coalescent simulations with different demographic scenarios (see *Supplementary Note* for the simulation protocols). We first tested whether MSMC can locally decode the time to the first coalescence. Figure 1b shows the posterior probability over the hidden state for two, four and eight haplotypes. As shown, the model recovers the true hidden state with good local resolution. As also seen in Figure 1b, the typical length of segments of constant hidden state increases with the sample size, while the typical time to the first coalescence decreases as described above.

We then tried to infer back the simulation parameters from the simulated sequence data, applying MSMC to two different demographic scenarios (Figure 2). First, we simulated a single population under a sequence of population growths and declines. As can be seen in Figure 2a, MSMC recovers the resulting zig-zag pattern with good resolution, where two haplotypes (similar to PSMC) yield good estimates between 50kya and 2,000kya, and eight haplotypes give estimates as recent as 2kya, with a small bias towards smaller population sizes in the more distant past We were able to test a reduced data set with reduced parameters with 16 haplotypes, as shown in *Supplementary Figure 2e*, which suggests that the bias observed with 8 haplotypes in the distant past increases further with more haplotypes (see Discussion). We also tested other simulated histories with sharp changes in population size, shown in *Supplementary Figures 2a,b*. As expected from the experience with PSMC, very rapid population size changes are smoothed out over an interval around the true change time.

**Figure 2:**
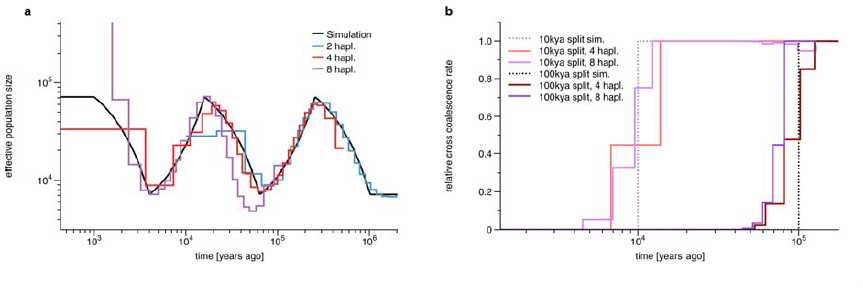
Testing MSMC on simulated data. (**a**) To test the resolution of MSMC applied to two, four and eight haplotypes, we simulated a series of exponential population growths and declines, each changing the population size by a factor 10. MSMC recovers the resulting zig-zag-pattern (on a double-logarithmic plot) in different times, depending on the number of haplotypes. With two haplotypes, MSMC infers the population history from 40kya to 3mya, with four haplotypes from 8kya to 300kya, and with eight haplotypes from 2kya to 50kya. (**b**) Model estimates from two simulated population splits 10kya and 100kya. The dashed lines plot the expected relative cross coalescence rate between the two populations before and after the splits. Maximum likelihood estimates are shown in red (four haplotypes) and purple (eight haplotypes). As expected, four haplotypes yield good estimates for the older split, while eight haplotypes give better estimates for the recent split.

Next, we simulated two population split scenarios, where a single ancestral population split into two equally sized populations 10kya and 100kya, respectively. For each population split scenario we inferred effective coalescence rates across the two populations and within populations, as a function of time (see *Online Methods*). We provide a measure for the genetic separation of populations as the ratio between the cross- and the within-coalescence rates, which we term the *relative cross coalescence rate.* This parameterization effectively models population separations and substructure in a simpler way than a standard forwards-in-time island-migration model, which would require a larger state space for structured genealogies, as shown in [10] (see also *Supplementary Note,* Chapter 10). Although not standard, we suggest that it is a more direct measurement of what can be derived about genetic exchange between historic populations from modern samples. The relative cross coalescence rate should be close to one when the two populations are well mixed, and zero after they have fully split. As shown in Figure 2b for four and eight haplotypes, our MSMC estimates correctly show this, although the instantaneous split time in the simulation is spread out over an interval around the split time. As expected, eight haplotypes yield better estimates for the more recent split scenario at 10kya, while four haplotypes yield better estimates for the older split at 100kya.

We also tested a population split with subsequent migration (*Supplementary Figure 2c*), for which we infer a higher relative cross coalescence rate across the two populations after the split, as expected. We further tested the robustness of our method under population size changes before and after the split (*Supplementary Figure 2d*), the consequences of the approximation to the singleton branch length (*Supplementary Note and Supplementary Figure 3*) and recombination rate heterogeneities (see *Supplementary Figure 4* and *Online Methods*), with no substantial effect on our estimates.

**Figure 3:**
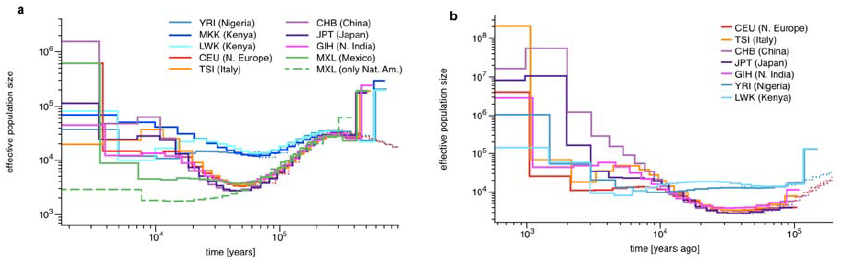
Population Size Inference from whole genome sequences. (**a**) Population size estimates from four haplotypes (two phased individuals) from each of 9 populations. The dashed line was generated from a reduced data set of only the Native American components of the MXL genomes. Estimates from two haplotypes for CEU and YRI are shown for comparison as dotted lines, (**b**) Population size estimates from eight haplotypes (four phased individuals) from the same populations as above except MXL and MKK. In contrast to four haplotypes, estimates are more recent. For comparison, we show the result from four haplotypes for CEU, CHB and YRI as dotted lines. Data for this Figure is available via *Supplementary Table 5*.

**Figure 4:**
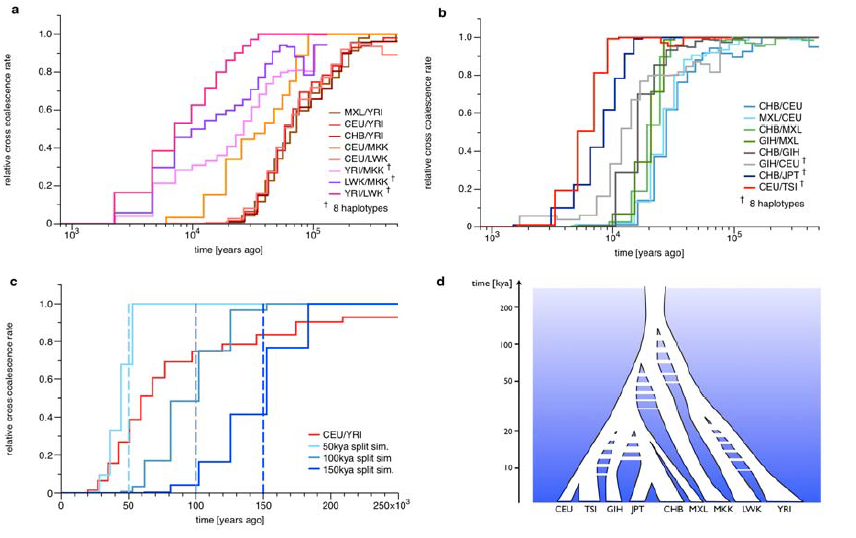
Genetic Separation between population pairs. (**a**) Relative cross coalescence rates in and out of Africa. African/Non-African pairs are shown in red colors, pairs within Africa in Purple colors, (**b**) Relative cross coalescence rates between populations outside Africa. European/East-Asian pairs in blue colors, Asian/MXL pairs in green colors, and other non-African pairs in other colors as indicated. The pairs that include MXL are masked to include only the putative Native American components. The most recent population separations are inferred from eight haplotypes, i.e. four haplotypes from each population, as indicated in the legend, (**c**) Comparison of the African/Non-African split with simulations of clean splits. We simulated three scenarios, at split times 50kya, 100kya and 150kya. The comparison demonstrates that the history of relative cross coalescence rate between African and Non-African ancestors is incompatible with a clean split model, and suggests it progressively decreased from beyond 150kya to approximately 50kya. (**d**) Schematic representation of population separations. Timings of splits, population separations, gene flow and bottlenecks are schematically shown along a logarithmic axis of time. Data for this Figure is available via *Supplementary Table 5*.

MSMC requires in principle fully phased haplotypes as input, although we can partially allow for unphased data at a subset of sites (see *Online Methods* for details). To test possibilities of extending our unphased approximation to entirely unphased samples, we simulated data sets with one or both individuals unphased. As shown in *Supplementary Figure 5*, population size estimates based on unphased data are still relatively accurate, although with biases at the two ends of the analyzed time range. Relative cross coalescence rate estimation based on partially or fully unphased data is less accurate and more biased in the distant past Because of this, when applying MSMC to real data we leave unphased sites in the population size estimate analyses, but remove them from the population split analyses (see also *Supplementary Figure 6*).

### Inference of population size from whole genome sequences

We applied our model to the genomes from one, two and four individuals sampled from each of 9 extended HapMap populations [20]: YRI (Nigeria), MKK (Kenya), LWK (Kenya), CEU (Northern European ancestry), TSI (Italy), GIH (North Indian ancestry), CHB (China), JPT (Japan), MXL (Mexican ancestry admixed to European ancestry) (details in *Supplementary Table 1*). We statistically phased all genomes using a reference panel (see *Online Methods*) and tested the impact of potential switch errors by comparing with family-trio-phased sequences which are available for CEU and YRI (see *Online Methods* and *Supplementary Figure 6*).

The results from two individuals (four haplotypes) are shown in Figure 3a. In all cases the inferred population history from four haplotypes matches the estimates from 2 haplotypes where their inference range overlaps (60kya-200kya, see thin lines for CEU and YRI in Figure 3, and *Supplementary Figure 7a*). We find that all non-African populations that we analyzed show a remarkably similar history of population decline from 200kya until about 50kya, consistent with a single non-African ancestral population that underwent a bottleneck at the time of the exodus from Africa around 40-60kya [21–23]. The prior separation of non-African and African ancestral population size estimates begins much earlier at 150-200kya, clearly preceding this bottleneck, as already observed using PSMC [7]. We will quantify this further below by directly estimating the relative cross coalescence rate over time. In contrast, we see only a mild bottleneck in the African population histories with an extended period of relatively constant population size more recently than 100kya. Between 30kya and 10kya we see similar expansions in population size for the CEU, TSI, GIH, and CHB populations. For the Mexican ancestors we see an extended period of low population size following the out of Africa bottleneck, with the lowest value around 15kya, which is particularly pronounced when filtering out genomic regions of recent European ancestry due to admixture (dashed line in Figure 3, see *Online Methods*). This extended bottleneck is consistent with estimates of the time that the Native American ancestors crossed the Bering Strait and moved into America [21, 24–26]. We repeated all analyses based on four haplotypes on a replicate data set, based on different individuals, available for all populations except MXL. All results are well reproduced and show differences only in the most recent time intervals, shown in *Supplementary Figure 8.*

Analyzing eight haplotypes from each population except from MXL and MKK (see *Online Methods*), we can see recent population size changes with higher resolution than with four haplotypes, as shown in Figure 3b. Results from eight haplotypes are compatible with those from four haplotypes beyond 10kya, with systematically slightly lower estimates in the range 10-30kya as expected from Figure 2a (*see Supplementary Figure 7b*). We get new insights more recently than 10kya, during the period of Neolithic expansion. Focusing first on Asian populations, we see that the ancestors of CHB increased rapidly from 10kya to effective population sizes of over a million 2kya. The GIH ancestors also increased early from around 10kya, a little more slowly than CHB, then leveled out around 4kya until recently. The JPT ancestral population appears to have split from CHB by 9kya and only increased slowly up until 3kya, since when it also increased very rapidly. In Europe, Northern European ancestors (CEU) experience a relatively constant effective population size between 2kya and 10kya, only increasing rapidly since 2kya; Southern European ancestors (TSI) have a consistently higher effective ancestral population size, appearing to show a more complex history of increase and decrease between 10kya and 3kya, and then increasing earlier than the CEU. As discussed previously [7], such a pattern of increase and decrease can result from admixture of previously separated populations, consistent with multiple waves of peopling of Europe with a substantial genetic component from earlier waves in Southern Europe [27]. In Africa, we see YRI (Yoruba) expanding earliest around 6kya, consistent with the introduction of agriculture and the Bantu expansion [28], followed by LWK (Luhya), in which prior to 6kya there is a long “hump” in ancestral population size extending back beyond 50kya, again possibly reflecting admixture within the last few thousand years between populations initially separated tens of thousands of years ago.

### Divergence from and within African populations

MSMC lets us explicitly study the genetic separation between two populations as a function of time by modeling the relationship of multiple haplotypes, half of which are from one population, and half from the other. As shown in Figure 4a, from analyzing four haplotypes for each pair of populations, we find that all relative cross coalescence rates between any non-African population and Yoruba are very similar and exhibit a slow gradual decline beginning earlier than 200kya and lasting until about 40kya. This similarity gives additional information beyond that from population size estimates, consistent with all non-African populations diverging as a single population from the Yoruban ancestors.

To understand whether the gradual decline in relative cross coalescence rate between YRI and non-African ancestors is due to the inability of MSMC to detect rapid changes (see Figure 2b) or due to true ongoing genetic exchange, we compared it with simulated clean split scenarios at three different time points in the past (see Figure 4c). This comparison reveals that no clean split can explain the inferred progressive decline of relative cross coalescence rate. In particular, the early beginning of the drop would be consistent with an initial formation of distinct populations prior to 150kya, while the late end of the decline would be consistent with a final split around 50kya. This suggests a long period of partial divergence with ongoing genetic exchange between Yoruban and Non-African ancestors that began beyond 150kya, with population structure within Africa, and lasted for over 100,000 years, with a median point around 60-80kya at which time there was still substantial genetic exchange, with half the coalescences between populations and half within (see Discussion). We also observe that the rate of genetic divergence is not uniform but can be roughly divided into two phases. First, up until about 100kya, the two populations separated more slowly, while after 100kya genetic exchange dropped faster. We note that the fact that the relative cross coalescence rate has not reached one even around 200kya (Figure 4c) may possibly be due to later admixture from archaic populations such as Neanderthals into the ancestors of CEU after their split from YRI [29].

In Figure 4a we also see extended divergence patterns based on eight haplotype analysis between the ancestors of the three African populations, with LWK and YRI closest, and MKK showing very slow increase in relative cross coalescence rate going back in time with YRI and LWK. These declines are all more gradual than those shown in Figure 4b between out-of-Africa populations, to be discussed further below, suggesting that they also are not clean splits but reflect gradual separation and perhaps complex ancestral structure with admixture. In addition, we see a different history between MKK and CEU compared to LWK and CEU, which in turn is very similar to the YRI/CEU estimates. Our results suggest that Maasai ancestors were well mixing with Non-African ancestors until about 80kya, much later than the YRI/Non-African separation. This is consistent with a model where Maasai ancestors and Non-African ancestors formed sister groups, which together separated from West African ancestors and stayed well mixing until much closer to the actual out-of-Africa migration. Non-zero estimates of relative cross coalescence rate between MKK and CEU after 50kya are probably confounded by more recent admixture from Non-African populations back into East African populations including Maasai [30, 31].

### Divergences outside Africa

As expected, the oldest split amongst out-of-Africa populations is between European and East Asian (CHB and MXL) populations, most of which occurs between 20-40kya (Figure 4b). Intriguingly there may be a small component (10% or less) of this separation extending much further back towards 100kya, not compatible with a single out-of-Africa event around 50kya. Next oldest is a separation between Asian (CHB and GIH) and American (MXL) populations around 20kya. This is the most rapid separation we see, compatible with a clean split. Passing over the GIH separations for now, we see from eight haplotype analyses separations between JPT and CHB around 8-9kya compatible with the divergence in population size history described above, and between CEU and TSI around 5-6kya, both also relatively sharp. *Supplementary Figure 9* shows four haplotype analyses of the same separations.

The pattern of divergence of the North Indian ancestors (GIH) from East Asian and European ancestors is more complex. We observe continued genetic exchange between GIH ancestors and both of these groups until about 15kya, suggesting that even though East Asians and Europeans separated earlier, there was contact between both of these and the ancestors of GIH after this separation, or equivalently that there was ancient admixture in the ancestry of Northern India. This deviation from a tree-like separation pattern is independently confirmed by D statistics from an ABBA-BABA test [32] (see *Online Methods* and *Supplementary Table 2*), which also indicate that GIH is genetically closer to CEU than to CHB and MXL. This is consistent with the slower and later decline in relative cross coalescence rate between CEU and GIH compared to between CEU and CHB (Figure 4b). These findings suggest that the GIH ancestors remained in close contact with CEU ancestors until about 10kya, but received some historic admixture component from East Asian populations, part of which is old enough to have occurred before the split of MXL.

## Discussion

We have presented here both a new method MSMC, and new insight into the demographic history of human populations as they separated across the globe. We have shown that MSMC can give accurate information about the time dependence of demographic processes within and between populations from a small number of individual genome sequences. Like PSMC it does this without requiring a simplified model with specific bottlenecks, hard population splits and fixed population sizes as do previous methods based on allele frequencies [33–35] or more general summary statistics [36–39]. However, MSMC extends PSMC to an order of magnitude more recent times and also allows us to explicitly model the history of genetic separations between populations. Because MSMC explicitly models the first coalescence between all possible pairs of haplotypes, it changes the time period considered in a quadratic fashion. This should be compared with the more naive approach of combining data from PSMC run on different individuals, which would increase information at most linearly, since individuals’ histories are not independent.

While MSMC substantially advances the methodology from PSMC to multiple samples and much more recent times we have also seen that its practical application appears to be limited to about eight haploid sequences, both because of the approximations involved as well as because of computational complexity (see *Supplementary Figure 2e* for estimates based on 16 haplotypes). It is intriguing, however, to imagine that larger numbers of samples in principle contain information about even more recent population history, potentially up until a few generations ago. The basic idea of looking at first coalescence events, as presented here, may lead to new developments that complement MSMC in this direction, e.g. by focusing on rare mutations such as doubletons in large samples [40].

The alternative conditional sampling approach mentioned in the introduction [9, 10], implemented in the software *diCal* also allows for higher resolution at more recent times. However, when we applied *diCal* to our zig-zag population size simulation it did not appear to infer population history more recently than about 20kya (see *Supplementary Figure 10*). Also there is currently no way to address the relationship between populations as done by MSMC as relative cross coalescence rate. Both of these points may improve in future developments of that method.

Applied to 34 whole genome sequences from nine populations, MSMC gives a picture of human demographic history within the last 200,000 years, beginning with the genetic separations of Yoruban from Non-African ancestors and extending well into the Neolithic (Figure 4d). We find strong evidence that the Yoruban/Non-African separation took place over a long time period of about 100,000 years, starting long before the known spatial dispersal into Eurasia around 50kya. Because we model directly an arbitrary history over time of relative cross coalescence rate between populations, we can see more clearly a progressive separation than earlier analyses based on a single separation time with some subsequent migration [7, 17, 33, 41]. However Yoruba does not represent all of Africa. We now see that the Maasai separation from the out-of-Africa populations occurred within the last 100,000 years. The older part of the separation from Yoruba may therefore be a consequence of ancient population structure within Africa, though the direct picture of relationships between African populations is complicated by extensive more recent exchange that we see between all three of Yoruba, Maasai and Luhya within the last 100,000 years. This scenario still does not rule out a possible contribution from an intermediate modern human population that dispersed out of Africa into the Middle-East or the Arabian peninsula but continued extensive genetic exchange with its African ancestors until about 50kya [17, 42, 43].

Our results are scaled to real times using a mutation rate of 1.25×10^−8^ per nucleotide per generation, as proposed recently [16] and supported by several direct mutation studies [14–16]. Using a value of 2.5×10^−8^ as was common previously [44, 45] would halve the times. This would bring the midpoint of the out-of-Africa separation to an uncomfortably recent 30-40kya, but more concerningly it would bring the separation of Native American ancestors (MXL) from East-Asian populations to 5-10kya, inconsistent with the paleontological record [25, 26]. We suggest that the establishment and spread of the Native American populations may provide a good time point for calibrating population genetic demographic models. We note that the extended period of divergence between African and non-African ancestors that we observe reconciles the timing of the most recent common ancestor of African and Non-African mtDNA around 70kya [1, 18] with the lower autosomal nuclear mutation rate used here, which in simple split models would suggest a separation around 90-130kya [7, 17, 33, 41, 46]. Given that we observe extensive cross-coalescence at nuclear loci around 60-80kya, sharing a common ancestor during that time for mtDNA, which acts as a single locus with reduced effective population size, is entirely likely.

## Acknowledgements

We thank Aylwyn Scally for useful comments and discussion, in particular on interpreting the population divergence estimates, and the Durbin group for general discussion. S.S. thanks Andrej Fischer for helpful support with the HMM implementation details. We thank Jeff Kidd, Simon Gravel and Carlos Bustamante for making ancestry tracts for the MXL individuals available to us. S.S. acknowledges grant support from an EMBO long-term fellowship. This work was funded by Wellcome Trust grant 098051.

## Author Contributions

R.D. proposed the basic strategy and designed the overall study. S.S. developed the theory, implemented the algorithm and obtained results. S.S. and R.D. analyzed the results and wrote the manuscript.

## Online Methods

### Sequence Data

Whole genome sequences were generated by Complete Genomics [11] and are freely available on their website (The “69 genomes” public data set). Homozygous and heterozygous consensus calls are taken directly from Complete Genomics’ tabular format file “masterVar” within the ASM directory of each sample. All regions with a quality tag less than “VQHIGH” were marked as missing data. Furthermore, sites for which more than 17 out of the 35 overlapping 35mers from the reference sequence can be mapped elsewhere with zero or one mismatch were marked as missing (this is similar to the filtering in [7]). The anonymous names of all individuals are summarized in *Supplementary Table 1.* The number of called sites and the diversity in each population is summarized in *Supplementary Table 3.* In the MKK samples, we detected cryptic relatedness (see *Supplementaiy Table 4*), by comparing the heterozygosity within and between samples. We find that this is due to a factor ¼ lower mutual heterozygosity between the two individuals NA21732 and NA21737. We therefore removed that population from the 8 haplotype analysis. We find that MXL individuals have a higher heterozygosity across than within samples, which may be due to different ancestry proportions.

### Phasing

All segregating sites that are present in the 1000 Genomes integrated variant call set were phased with the SHAPEIT2 software [47] using the 1000 Genomes reference panel [48]. In addition we generated full chromosomal haplotype phases from available trio data for YRI and CEU to assess the impact of the accuracy of the population phasing method vs. trio phasing. Current switch error rates when phasing against a 1000 Genomes reference panel are estimated to be about 500kb-1Mb (J. Marchini, personal communication), which limits the possibility to detect IBD-segments to times older than a few thousand years ago, which is at the lower limit of our analysis based on eight haplotypes.

As shown in *Supplementary Figure 5*, unphased sites affect population size estimates and relative cross coalescence rate estimates differently. From a direct comparison with trio-phased data we find that population size estimates are better if unphased sites are included in the analysis (taken as multiple observations in the HMM, see below) while for relative cross coalescence rate estimates, unphased sites generate severe artifacts in recent time intervals (see *Supplementary Figure 6*). The reason for this is the high sensitivity on shared long haplotypes between individuals, which are not detected if unphased sites are scattered across the sequence. We therefore removed unphased sites for the population split analysis, which in particular improves recent population divergence estimates after 50kya. In order to remove unphased sites in an unbiased way, we also removed all blocks of homozygous calls which end in an unphased heterozygous site, marking them as missing data. This is necessary because otherwise the removal of unphased sites would generate long segments of homozygous calls which are interpreted as long IBD segments of very recent tMRCA.

### Admixture masks for MXL

Admixture masks for NA19735 were downloaded from the 1000 Genomes project website as part of the phase I analysis results. Admixture tracts for the remaining MXL individuals were generated by J. Kidd and Coworkers [49] and kindly made available to us. For our population split analysis with four haplotypes we only used the two individual with the highest Native American component as MXL representative, NA19735 and NA19670. We then filtered out all segments in the genome that were not annotated as homozygous Native American.

### D Statistics (ABBA-BABA test)

We used the following samples: NA20845 (GIH), NA12891 (CEU), NA18526 (CHB) NA19735 (MXL), NA19238 (YRI). For the D statistic, we count sites at which YRI is homozygous and define the respective allele as “ancestral” (A). We then require that the individual specified in the third column carries at least one derived allele (B). We also require that of the individuals specified in the first and second column one is homozygous for allele A, and the other carries at least one derived allele (B). This leaves two configurations per setup named ABBA and BABA, respectively, see *Supplementary Table 2.* The D-statistic is then: D = (#(ABBA) – #(BABA)) / (#(ABBA) + #(BABA)). We compute p-values based on a two-tailed binomial test (see *Supplementary Table 2*). We performed this test for the two triples CEU/GIH/CHB and CEU/GIH/MXL, using YRI as outgroup. For the reversed scenarios, treating CEU as donor population and CHB or MXL as first population in the test, D scores are higher, suggesting that GIH is genetically closer to CEU than to CHB and MXL.

### MSMC Model

The Multiple Sequential Markovian Coalescent (MSMC) is a Hidden Markov model along multiple phased haplotypes. Its hidden state is a triple (*t, i, j*), where *t* is the first coalescence time of any two lineages, and *i* and *j* are the labels of the two lineages participating in the first coalescence. As detailed in *Supplementary Note,* we need additional local information about the genealogical tree, namely the singleton branch length *T_s_*, which sums the lengths of all branches that give rise to variants of minor allele count 1 if mutations occur on these branches. The singleton branch length can be estimated in a separate step before the main inference. We neglect the dependency of the transition rate on the singleton branch length, which is an approximation that is validated by simulations, see *Supplementary Figure 3.*

### Transition Rate

We use the Sequential Markovian Coalescent framework [5, 6, 50], SMC’, to derive transition rates between hidden states. As shown in *Supplementary Note* with help of additional illustrations, the transition probability to state (*t, i, j*) from state (*s, k, l*) can be derived relatively straightforwardly, and separated into the probability that no change of the first coalescence time occurs, either because no recombination event happened, or because the recombination event does not affect the first coalescence:

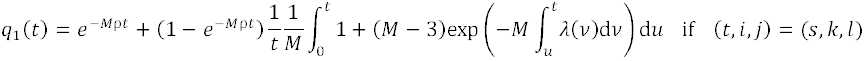

and the probability to change from time *s* to time *t* due to a recombination event:

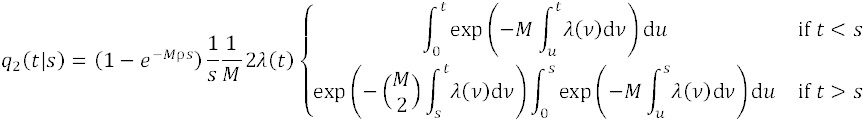

Here, M denotes the number of haplotypes, ρ is the scaled recombination rate and λ(t) is the scaled inverse population size as a function of time. When samples from different subpopulations are analyzed, the transition rate can be modified to allow for different coalescence rates within and across populations, see *Supplementary Note.*

For only two haplotypes, MSMC is very similar to PSMC [7], with subtle differences due to the different underlying model SMC’ [6] versus SMC [5]. We therefore call our special case of two haplotypes PSMC’ to distinguish it from PSMC.

### Emission Rate

For the emission rate we assume that we are given local estimates of the singleton branch length of the tree, *T_s_* (see *Supplementary Note*), which is treated as a “soft” constraint, that is we always consider max (*T_s_, Mt*) when we write *T_s_* For more than 2 haplotypes, an observation of alleles given some state (*t, i, j*) can be classified into the following categories, each of which has its own emission probability *e*(*t; T*_s_), derived in *Supplementary Note:*

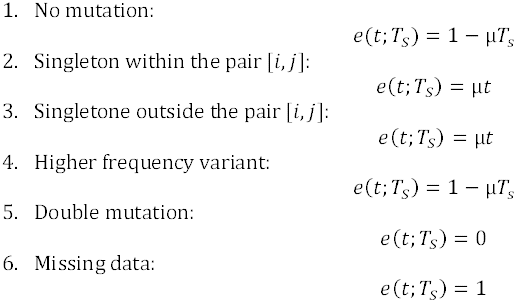

If sites have multiple observations from ambiguous phasing results, we average the emission probability over those observations.

### Parameter Inference

To infer parameters with MSMC, we discretize time into segments, following the quantiles of the exponential distribution. The boundaries of the segments are

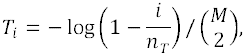

where *n_T_* is the number of segments, and *M* is the number of haplotypes, which determines the expected time to first coalescence. All times are given in units of 2*N*_0_ generations, where 2*N*_0_ is fixed from Watterson’s estimator. The number of segments used in the population size analysis was *n_T_ =* 40, and in the estimation of relative cross coalescence rate *n_T_* = 30. The denominator reflects the quadratically decreased time to first coalescence with increased sample size M. The limits on the inference are given by the second and the second-last boundary, which correspond to the 2.5%- and the 97.5%-quantile boundary of the distribution of first coalescence times.

As in PSMC, we join segments to reduce the parameter search space. The segmentation we used was 10*1+15*2 for the population size inference, i.e. the 30 rightmost segments where joined to pairs of 2. For the relative cross coalescence rate estimates we used a pattern of 8*1+11*2. For plots on a logarithmic time axis we chose the left cutoff to be at *T*_1_/4 and omitted the last time interval, as its value is shown also by the joined second last time interval.

In each segment, the coalescence rate is kept constant. If all samples are from the same population, we have one parameter per segment, which is the scaled inverse population size, λ_i_ where *i* enumerates the segments in time. If samples from two populations are analyzed, we have three parameters per segment: λ^1^_i_, λ^12^_i_, and λ^2^_i_. Here, λ^1^_i_ and λ^2^_i_ denote the coalescence rates within the two populations, and λ^12^_i_ denotes the coalescence rate across populations. In principle, there are two further free parameters: the scaled recombination rate ρ and the scaled mutation rate θ. In practice, we fix θ from Watterson’s estimator for θ. The recombination rate can be inferred relatively well from PSMC’, see *Supplementary Figure 1*, and we fix it to that estimate for more than two haplotypes to reduce the parameter search space. We note that inference results on the coalescence rates are relatively independent of the recombination rate.

Maximum likelihood estimates of all free parameters are generated iteratively by means of the Baum-Welch algorithm [51, 52] with a coarse-grained Q-function and numerical maximizations using Powell’s Direction-set method. For the coarse-graining of the Q-function, we use precomputed transition-emission matrices and evaluate the local variables only every 1000bp. This optimization makes it possible to analyze whole human genomes without the need to first bin the data into windows of 100bp, as in [7].

For only one population analyzed, an estimate of the scaled population size in time interval *i* is directly given as the inverse of the coalescence rate: N_i_/N_0_=l/λ_i_. The scaling parameter N_0_ is fixed from the scaled mutation rate (see above): N_0_= θ/(4μ), where μ=1.25x10^−8^ is the per site per generation mutation rate. Relative cross coalescence rate estimates are obtained by dividing the cross-population coalescence rate by the average within-population coalescence rate: γ_i_ = 2λ^12^_i_ / (λ^1^_i_ + λ^2^_i_). In the maximization step of the Baum-Welch algorithm we constrained the optimization to γ_i_≤1.

### Implementation

We implemented the model and inference algorithm in the D programming language (see URL below). The source code together with executables for common platforms is freely available from Github (see URL below). We also provide additional information on the input file format and necessary pre-processing in the *Supplementary Note.*

## URLs

D Programming language: http://www.dlang.org

Github: https://github.com/stschiff/msmc

